# PTPRU, a quiescence-induced receptor tyrosine phosphatase negatively regulates osteogenic differentiation of human mesenchymal stem cells

**DOI:** 10.1101/2021.04.29.442062

**Authors:** Mohammad Rumman, Jyotsna Dhawan

## Abstract

Bone marrow mesenchymal stem cells (MSCs) are heterogeneous osteo-progenitors that are mainly responsible for bone regeneration and homeostasis. In vivo, a subpopulation of bone marrow MSCs persists in a quiescent state, providing a source of new cells for repair. Previously, we reported that induction of quiescence in hMSCs *in vitro* skews their differentiation potential in favour of osteogenesis while suppressing adipogenesis. Here, we uncover a new role for a protein tyrosine phosphatase, receptor type U (PTPRU) in repressing osteogenesis during quiescence. A 75 kD PTPRU protein isoform was found to be specifically induced during quiescence and down-regulated during cell cycle reactivation. Using siRNA-mediated knockdown, we report that in proliferating hMSC, PTPRU preserves self-renewal, while in quiescent hMSC, PTPRU not only maintains reversibility of cell cycle arrest but also suppresses expression of osteogenic lineage genes. Knockdown of PTPRU in proliferating or quiescent hMSC de-represses osteogenic markers, and enhances induced osteogenic differentiation. We also show that PTPRU positively regulates a β-catenin-TCF transcriptional reporter. Taken together, our study suggests a role for a quiescence-induced 75kD PTPRU isoform in modulating bone differentiation in hMSC, potentially involving the Wnt pathway.

## Introduction

Adult stem cells (ASC) that reside in normal uninjured tissues persist in a quiescent or G0 state. Quiescence is a reversible cell cycle arrest that is triggered in response to growth-inhibiting signals, or by the absence of growth-promoting signals (Coller et al. 2006). In response to tissue injury and damage, quiescent ASCs are activated by secreted or mechanical signals arising from the injured tissues. ASCs proliferate and differentiate to repair the damaged tissue (Schultz et al. 1978, Cotsarelis et al. 1990, Cheshier et al. 1999). Importantly, a few activated SCs return to quiescence and restore the resident tissue stem cell pool. Quiescence contributes to maintenance of the steady state number of ASCs available in adult tissues, as well as plays a role in protecting cells against proliferation-associated stresses (Arai et al. 2004, Rossi et al. 2007, Mourikis et al. 2012), and the activation of alternate programs such as senescence and apoptosis. Thus, quiescence is a key feature of ASC function, and loss of quiescence in aging and pathological conditions is associated with compromised regeneration and tissue function (Cheng et al. 2000, Chakkalakal et al. 2012, Garcia-Prat et al. 2016).

Quiescence is characterized by induction of Snf and PKC pathways and suppression of PKA and mTOR pathways (Gray et al. 2004), which is conserved from yeast to mammals (Dhawan and Laxman 2015). Growth factor signaling pathways such as PKA are essential for proliferation, and the decline of proliferation-inducing signals are well established to result in quiescence (Thevelein and de Winde 1999, Gray et al. 2004). Quiescence-specific cross-talk of ubiquitous pathways also leads to re-wiring of signaling circuits resulting in G0-specific transcriptional outcomes (Aloysius et al. 2018). However, signaling pathways that actively promote or maintain quiescence, are less well understood, particularly in human stem cells (Rumman et al. 2015).

MSCs are a heterogeneous population of stem cells that have the ability to self-renew and to predominantly differentiate into cells of mesodermal lineages such as adipocytes, osteocytes, and chondrocytes in vitro (Pittenger et al. 1999, Jiang et al. 2002), with some evidence for trans-differentiation into neuronal and epithelial cells (Barui et al. 2018, Urrutia et al. 2019). However, *in vivo*, MSCs primarily contribute to bone development and repair (Kassem et al. 2004, Kassem and Bianco 2015). Besides their ability to serve in replacement of damaged tissue, increasing evidence indicates that MSCs also possess immuno-modulatory and anti-inflammatory properties via secretion of a variety of cytokines and growth factors (Le Blanc and Ringden 2007, Petrie Aronin and Tuan 2010). Thus, MSCs are ideal candidates for tissue engineering, regenerative medicine and treatment of autoimmune disease. A growing body of evidence suggests that a subpopulation of endogenous MSCs enter quiescence in vivo (Peiffer et al. 2007). However, the difficulty in isolating quiescent MSCs has posed a challenge in understanding the regulation of this reversibly arrested state.

Previously, we showed that hMSC can be induced to enter a reversibly arrested state when cultured in non-adherent conditions, despite the presence of growth-promoting concentrations of serum (Rumman et al. 2018). As seen in other mesenchymal cell types such as fibroblasts and myoblasts (Dike and Farmer 1988, Dhawan and Helfman 2004, Sellathurai et al. 2013), quiescent hMSC will re-enter proliferation as synchronized populations when replated on an adhesive surface. These post-quiescent hMSC show enhanced stem cell properties such as self-renewal, and display an altered transcriptome. Notably, post-quiescence, these cells display an altered cell fate potential with increased osteogenic differentiation and reduced adipogenic differentiation (Rumman et al. 2018), mimicking their function in vivo. Thus, induction of quiescence in hMSCs in vitro enhances clinically relevant properties, and suggests that quiescent MSCs might be preferable to asynchronously proliferating MSCs as cell replacement therapy in bone fractures.

In this study, we report a new quiescence-induced regulator of osteogenic differentiation in hMSCs, the receptor-type protein tyrosine phosphatase U (PTPRU). PTPRU has been reported to possess two pseudo-phosphatase domains, but despite the absence of catalytic activity, PTPRU can recruit substrates, acting as a scaffold to compete for binding to protein substrates (Hay et al. 2020). Herein, we analyzed the role of PTPRU in mediating the fate-changes seen in hMSC quiescence, and find that this receptor-type phosphatase plays a role in mitigating osteogenic differentiation in G0, potentially via effects on Wnt/β-catenin signaling. Since induction of overt differentiation during arrest leads to a loss of stem cell self-renewal, we conclude that PTPRU plays a role in maintenance of reversibility of quiescence in hMSC.

## Materials and Methods

### Ethics Statement

Ethical approval for experimental studies on human stem cells was obtained from the InStem Institutional Ethics Committee and Institutional Committee for Stem Cell Research.

### Cell culture

Human bone marrow MSC were purchased from Texas A&M Health Science Centre, College of Medicine, Institute for Regenerative Medicine, Texas (Smith et al. 2004). Purchased hMSC were derived from male healthy donors aged between 20-30 years. hMSC were routinely cultured in MEM-alpha medium (Gibco#A10490-01) supplemented with 16% FBS (Gibco#16000) and 100 U/ml Pen-Strep (Gibco#15140-122) at 37 °C under humidified conditions. Cells were passaged regularly at 60-70% confluency. To minimize variability and to maximize potency, hMSC between passages 1-3 only were used in the study.

### Osteogenic and adipogenic differentiation of hMSC

hMSC were induced to osteogenic and adipogenic lineage as described previously (Pittenger et al. 1999, Rumman et al. 2018). Briefly, for osteogenesis and adipogenesis, hMSC were seeded at a density of 5000 cells/cm^2^. Osteogenic induction was initiated when cultures were at 65-70% confluency, using osteogenic differentiation medium (growth medium supplemented with Dexamethasone (10 nM) (Sigma#D4902), β-glycerolphosphate (20 mM) (Sigma#G9422), L-Ascorbic acid 2-phosphate (50 μM) (Sigma#49752)). At the end of 21 days, differentiated osteoblasts were stained with Alizarin Red S to visualize calcium deposition. To quantify calcium deposition, stained Alizarin Red S was eluted from cultures using cetylpyridinium chloride (Sigma#C0732) (Rumman et al. 2018) and absorbance measured at 405 nm. The stained undifferentiated control was used as blank. Adipogenesis was induced at 80-85% confluency using adipogenic differentiation medium (growth medium supplemented with Dexamethasone (0.5 μM) (Sigma#D4902), Isobutylmethyl-xanthine (0.5 μM) (Sigma#I7018) and Indomethacin (50 μM) (Sigma#I7378)) (Pittenger et al. 1999).

### Colony Forming Unit-fibroblast (CFU-f) assay

CFU-f assay was performed as described previously (Friedenstein et al. 1976). Briefly, cells were counted using Countess Automated Cell Counter (Life Technologies) and 500 viable cells per condition were plated in 100 mm dishes in growth medium. After 10 days, media was removed, and the dish was washed with PBS and colonies were stained with 2% methylene blue solution (in absolute ethanol (w/v)) for 15 mins at room temperature. Methylene blue was removed and dishes were washed under running water to remove any residual methylene blue. Colonies were counted manually using a stereomicroscope. Three replicate plates per condition were analyzed.

### BrdU incorporation assay

hMSC were incubated with 10 μM BrdU in growth medium for one hour. Cells were washed with PBS thrice and fixed in 4% PFA for 15 min at room temperature. Fixed cells were washed thrice with PBS, treated with 2N HCL in PBS-T for 30 min at 37 °C for DNA denaturation and washed with NaBH4 solution thrice to neutralize HCL. Blocking was done with 10% FBS in PBS-T at room temperature for 1 h. Cells were stained with anti-BrdU antibody (DSHB #G3G4) for 1 h at room temperature followed by incubation with secondary antibody (anti-mouse, Alexa Fluor 488) for 1 h at room temperature. Cells were again washed with PBS three times, and mounted using Vectashield Antifade Mounting Medium with DAPI (Vector laboratories #H-1200). 200-300 nuclei were counted for each sample, under Zeiss Axiovert 40 CFL microscope.

### Methyl cellulose preparation and suspension culture

2% methyl cellulose (MC) medium was prepared as described previously (Rumman et al. 2018). MSC were cultured in methyl cellulose (1.3% final concentration) at a density of 1 million cells per 10 ml suspension culture. Cells were harvested at designated time points by diluting MC with warm PBS, and replated at 1 million cells per 100 mm dish for RNA/protein.

### RNA isolation and qPCR

RNA was isolated using TRIzol reagent according to manufacturer’s recommendation. qPCR was performed on ABI 7900HT machine. Data was normalized using GAPDH as described (Rumman et al. 2018).

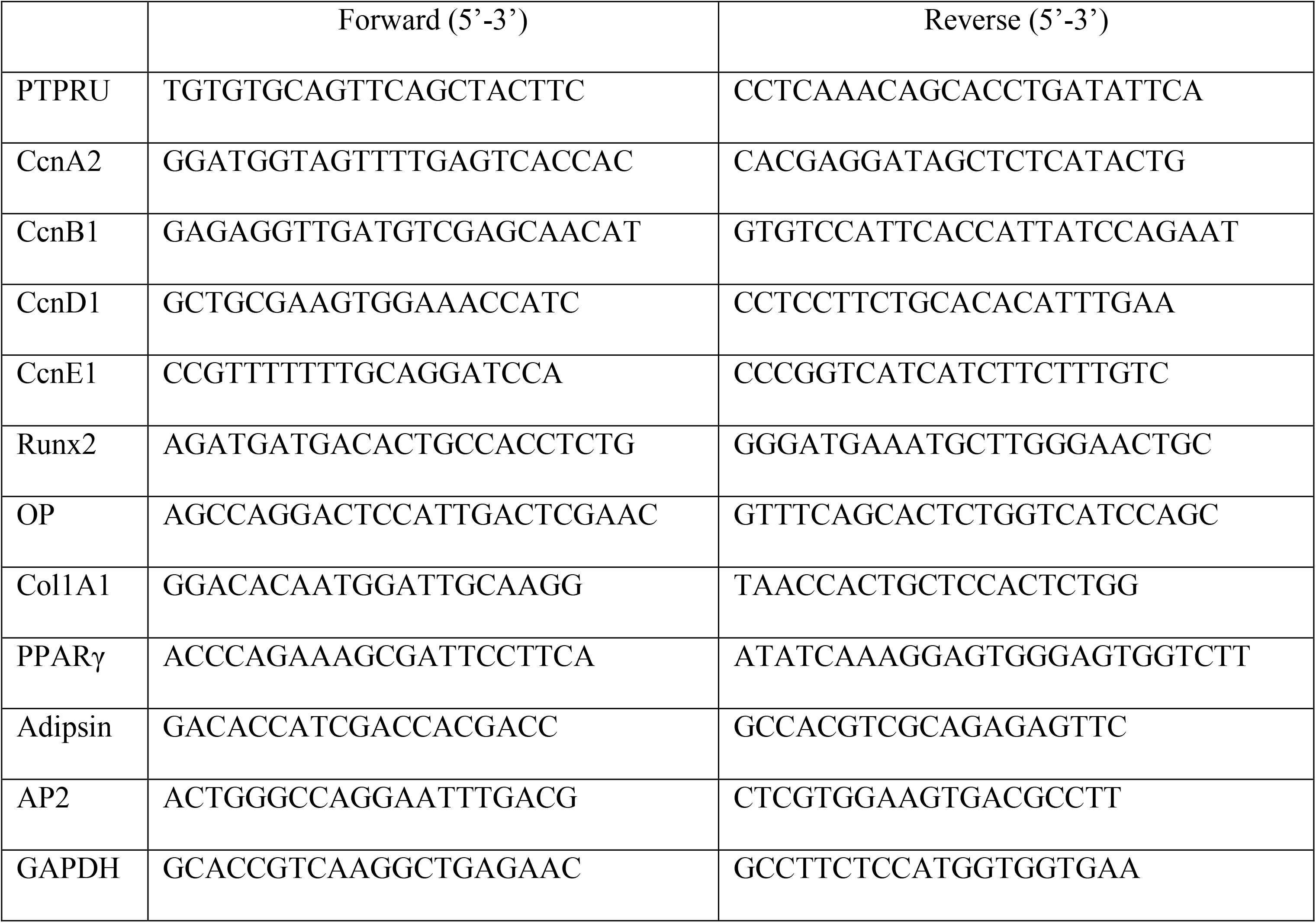

### siRNA transfection

siRNA against PTPRU (a pool of multiple target-specific siRNAs) was used for knockdown (Santa Cruz #sc-62910). For transfection in 6-well dishes, cells were plated at a confluency of 60-70% in growth medium. After 12 h MSCs were transfected using Lipofectamine RNAiMAX transfection reagent (Invitrogen#13778). Per well, 60 picomoles siRNA and 11 μl of RNAiMAX were diluted separately in 250 μl of Opti-MEM (Gibco#11058-021). These diluted solutions were mixed together, incubated at room temperature for 15 min, and the mixture added to cells for 24 h of transfection.

### TOP-flash Luciferase reporter assay

Adherent hMSC were transfected with PTPRU siRNA or scrambled (control, non-targeting) siRNA. After 12 h cells were super-transfected with the TCF reporter Top-flash (Subramaniam et al. 2013) along with Renilla-luciferase for normalization. 12 h later cells were suspended in MC, harvested for 48 h, and luciferase assay performed using a dual luciferase chemiluminescence detection kit (Promega #E1910) according to the manufacturer’s protocol. Around 5000 cells were lysed using 100 μl 1X lysis buffer and spun at 8000g for 10 min at 4 °C to pellet cell debris. 50 μl of the lysate was used for the detection of luminescence in a Sirius single tube luminometer. The normalized luciferase activity is represented as Relative Luminescent Units (RLU/μg protein) in bar diagrams.

### Statistical analysis

All experiments were repeated three times (3 biological replicates, each containing 3 technical replicates as needed) and standard deviation was calculated, significance of change compared to control was calculated by paired Student’s T-test (two tailed). Values showing ***p***-value less than 0.05 are marked by asterisk (*).

## Results

### Expression of PTPRU is restricted to reversibly arrest hMSC

Earlier, we reported the global transcriptomic changes that accompany entry of hTERT-MSC into reversible quiescence (G0) vs. differentiation into osteoblasts or adipocytes (Rumman et al. 2018). We found that several classes of phosphatases were specifically and reversibly induced in quiescent state. Of these, the receptor type protein tyrosine phosphatase PTPRU was substantially induced at the transcript level (Fig 1a). In order to validate the expression pattern of PTPRU, we used qRT-PCR analysis to detect its transcript levels in proliferating, quiescent and differentiated hMSC. Primers used in qRT-PCR targeted the exon 3 of PTPRU mRNA, which corresponds to the MAM (<u>M</u>eprin, <u>A</u>-5 protein, and receptor protein-tyrosine phosphatase <u>m</u>u)-domain of the protein (Fig 1b). We found that PTPRU transcript is detected in proliferating hMSC, but its expression is up-regulated ~200-fold in quiescent hMSC, returning to low levels when G0 cells were reactivated for 24 hrs (Fig 1c). Further, PTPRU expression in G0 was found to be even higher than in terminally differentiated osteoblasts and adipocytes (Fig 1c). This expression pattern indicates that transcriptional induction of PTPRU is specific to reversible arrest (G0) rather than terminal arrest that accompanies differentiation. PTPRU transcript is predicted to encode a protein of ~200 kD (Yan et al., 2006). Smaller fragments of PTPRU (~130 kD, ~80 kD and ~55 kD) have also been reported in a variety of cancer cell lines (Zhu et al. 2014). A commercially available antibody generated against full-length recombinant human PTPRU protein reproducibly detected a single band at ~75 kD which showed reversible induction in quiescent hMSC (Fig 1d). Neither predicted full-length PTPRU (200 kD), nor other reported short fragments (130 kD & 55 kD) were detected in hMSC (Fig 1d). The induction of ~75 kD PTPRU isoform in quiescent hMSC was reversible, as its expression was down-regulated upon reactivation of hMSC into the cell cycle.

**Fig 1.**
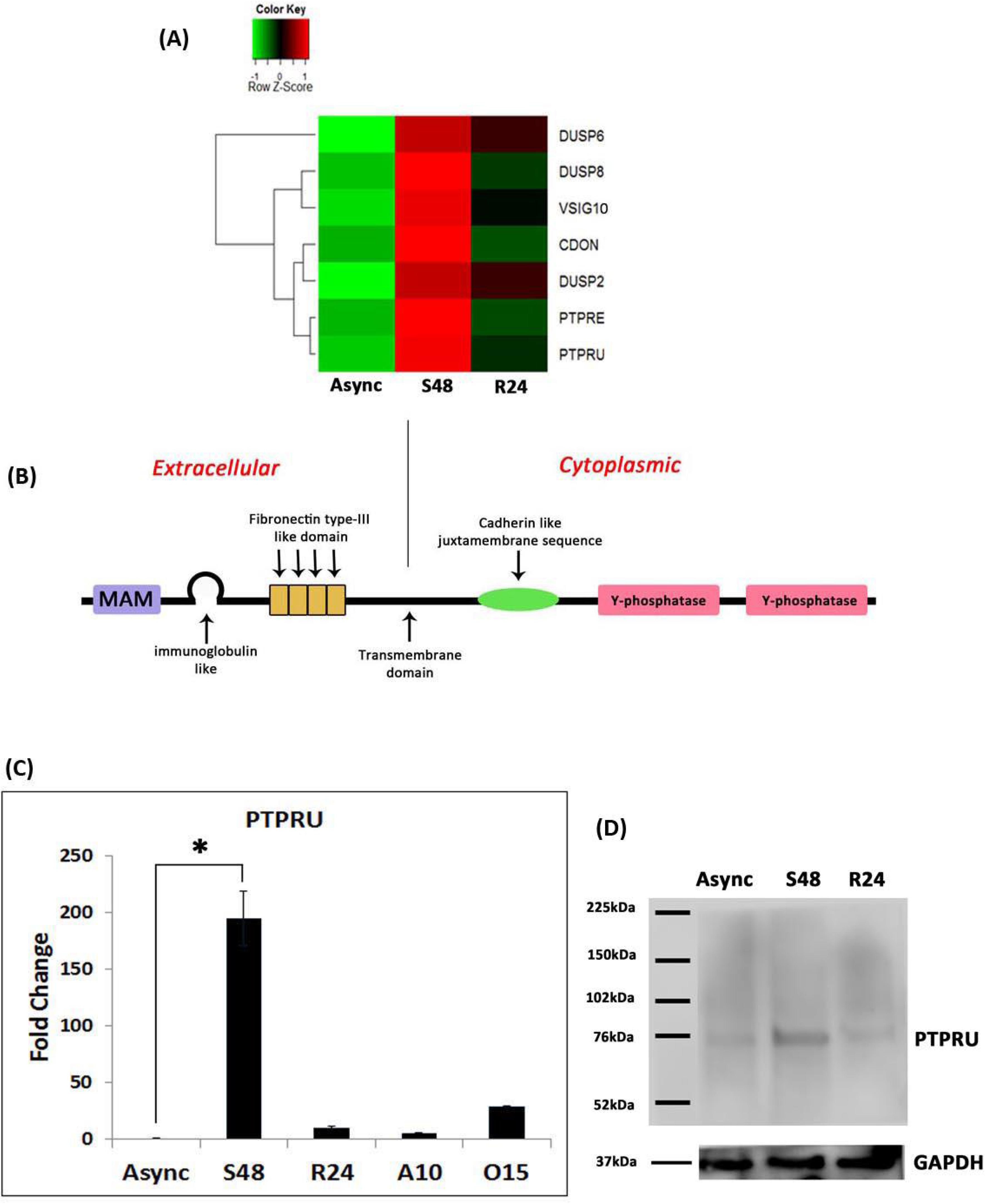
PTPRU expression in quiescent MSC. (A) Heatmap of transcripts encoding various phosphatases induced in quiescent hMSC. Data represents fold changes in transcript levels taken from microarray analysis (Rumman et al, 2018) and can be retrieved at GEO Accession # GSE60608. Asynchronously proliferating hMSC (Async), hMSC cultured in suspension for 48 hours to achieve quiescence (S48), reactivation of quiescent hMSC for 24 hours (R24). (B) Schematic showing domain architecture of full length PTPRU trans-membrane protein. (C) Transcript level of *PTPRU* in proliferating (Async), quiescent (S48), reactivated MSC, 24 hours post-quiescence (R24), hMSC differentiated for 10 days in adipogenic media (A10), hMSC differentiated for 15 days in osteogenic media (O15). Values represent mean ± S.D, n = 3, p< 0.05. (D) Expression of a single 75 kD isoform of PTPRU in asynchronous, quiescent, and reactivated MSC. GAPDH is shown as a loading control. Data are representative of 2 independent experiments

### Suppression of PTPRU in MSC slows proliferation

As PTPRU was found to be expressed in proliferating hMSC and was upregulated during quiescence, we used siRNA-mediated knockdown to elucidate the role of this phosphatase in both proliferating and quiescent conditions. Reduction of PTPRU mRNA levels was confirmed by qRT-PCR and western blot 24 hours after siRNA transfection. Greater than 90% knockdown was achieved at both RNA and at protein levels (Fig 2a).

**Fig 2.**
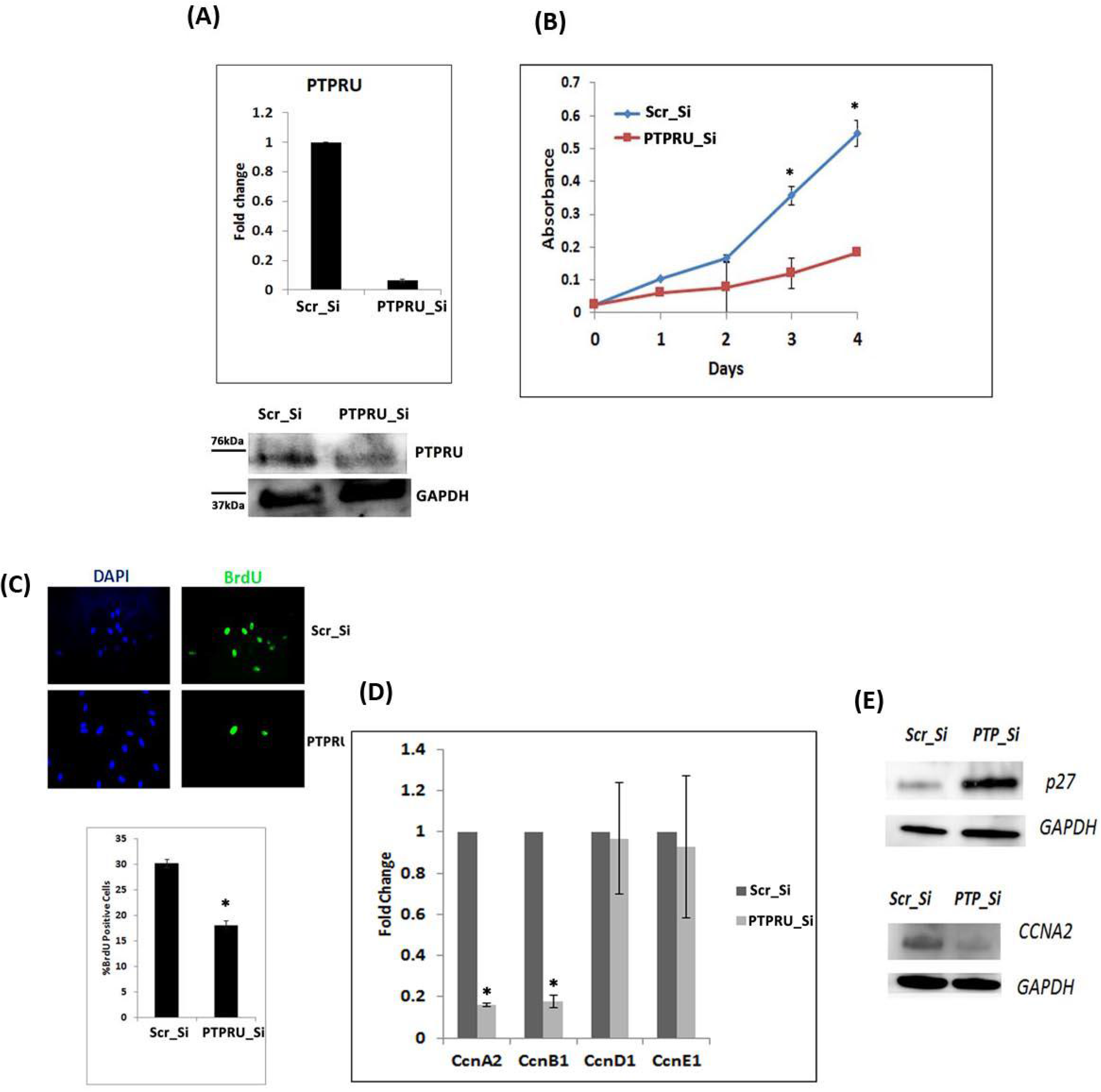
PTPRU knockdown in adherent proliferating MSC leads to altered cell cycle regulator expression. (A) siRNA-mediated knockdown of PTPRU in adherent proliferating MSC after 24 h of transfection: mRNA levels measured by qRT-PCR (upper panel) and protein levels by immunoblotting (lower panel). GAPDH is used as a loading control. (B) WST assay to assess rate of proliferation in PTPRU knockdown proliferating MSC. [mean + S.D, n=3, p< 0.05] (C) BrdU incorporation in PTPRU KD MSC in adherent condition. [mean + S.D, n=3, p< 0.05] (D) Transcript levels of *Ccna2, Ccnb1, Ccnd1* and *Ccne1* in PTPRU KD MSC in proliferating condition (E) Protein level of P27 and CCNA2 in PTPRU KD adherent MSC.

Since PTPRU expression was upregulated ~200-fold during quiescence and suppressed during the return to proliferation, we hypothesised that RNAi-mediated suppression of this gene may result in more rapid proliferation of MSC and/or failure of MSC to undergo cell cycle arrest doing suspension culture. Therefore, we assessed the rate of cell proliferation and the fraction of cells entering S-phase. Contrary to our hypothesis, in adherent proliferating hMSC, PTPRU knockdown resulted in slower proliferation as compared to control cells (Fig 2b). This observation was confirmed by BrdU incorporation assay, where knockdown cells in adherent condition showed ~2-fold less BrdU incorporation as compared to control cells (Fig 2c). Consistent with these findings, PTPRU knockdown cells showed repressed levels of cyclin A2 and cyclin B1 (Fig 2d), and elevated levels of cyclin dependent kinase inhibitor p27 (Fig 2e). Together, these results suggest that PTPRU is involved in maintaining the cell cycle in adherent proliferating cultures.

### hMSC lacking PTPRU arrest efficiently in G0, but exhibit altered expression of cell cycle regulators

To further dissect the role of PTPRU in MSC, we assessed the consequence of knockdown on the ability to enter quiescence. As MSC cultured in methylcellulose cannot be efficiently transfected, we transfected adherent MSC with anti-PTPRU siRNA, and 12 hours post transfection, trypsinized and suspended the cells in methylcellulose. After 2 days of culture, the cells were harvested and analyzed for PTPRU expression, BrdU incorporation and other markers. Cells harvested from suspension culture showed greater than 80% knockdown of PTPRU at 0 h and 24 h of reactivation post quiescence (Fig 3a,b). In suspension culture, PTPRU knockdown cells showed no increase in BrdU incorporation indicating that PTPRU is not required to either enter or maintain the arrested state (Fig 3d).

**Fig 3.**
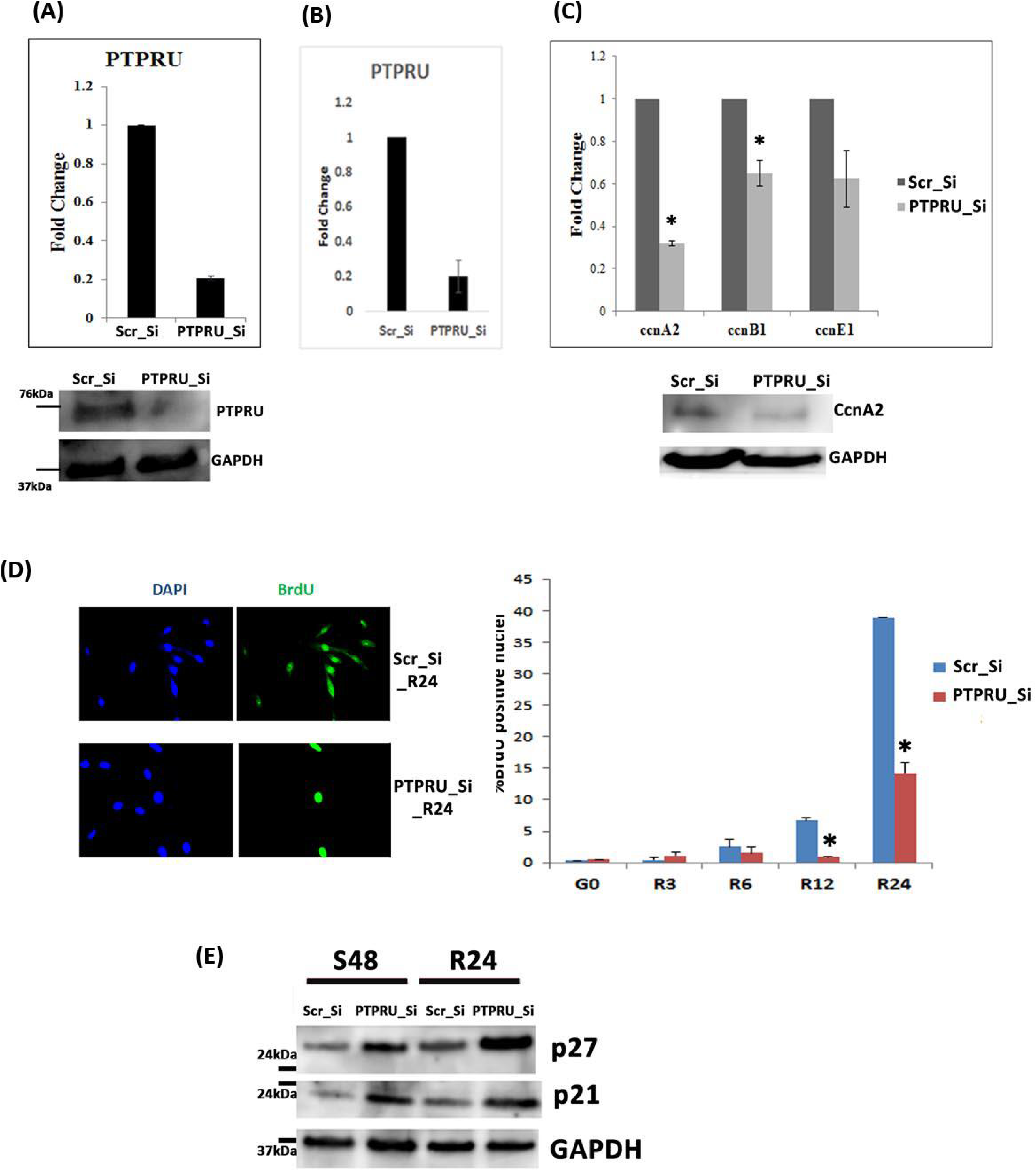
PTPRU knockdown during suspension and replating leads to altered hMSC quiescence and compromises reactivation. (A) siRNA-mediated knockdown of PTPRU during quiescence (mRNA and protein levels) (B) PTPRU knockdown in reactivated MSC after 24 h (mRNA levels) (C) Upper panel: Cell cycle transcript levels of *Ccna2, Ccnb1, and Ccne1* are down-regulated in PTPRU knockdown hMSC in quiescence; Lower panel: CCNA2 protein levels are reduced. (D) BrdU incorporation in PTPRU knockdown hMSC is reduced during reactivation; left panel shows a representative immuno-fluorescence image, right panel shows quantitation of BrdU incorporation over a replating time course of 3 to 24 hours (R3-R24). (E) Protein levels of the CDKIs P21 and P27 are increased in PTPRU KD MSC both during quiescence (S48) and reactivation (R24).

To determine whether the growth-arrested state achieved in PTPRU knockdown cells in suspension was molecularly similar to that of control G0 cells, we analyzed the expression of cell cycle regulators. We found that cyclin A2 mRNA was further suppressed and cell cycle inhibitors p21 and p27 were over-expressed in PTPRU knockdown cells as compared to control quiescent cells (Fig 3c,e). This observation suggests that PTPRU knockdown hMSC do arrest in suspension, but hyper-repression of cyclins and increased expression of CDKIs might affect their re-entry into cell cycle.

### hMSC lacking PTPRU are defective in cell cycle re-entry from G0

To assess whether PTPRU affects cell cycle re-entry following arrest, we replated suspension-arrested knockdown cells and measured BrdU incorporation. PTPRU knockdown was sustained at 24 hours post reactivation as confirmed by qRT-PCR (Fig 3b). PTPRU knockdown MSC show delayed and blunted reactivation kinetics: 12 hours after replating, PTRPU knockdown cells showed ~3-fold less BrdU positive cells when compared to control cells and ~2.5-fold less at 24 hours (Fig 3d). Failure of knockdown cells to reactivate efficiently was further supported by the observation that of the cell cycle inhibitors p21 and p27 continued to be expressed at high levels in reactivated knockdown cells (Fig 3e), compromising their ability to re-enter cell cycle efficiently. Further, loss of PTPRU caused loss of self-renewal in adherent MSC (Fig 4a) and post-G0 MSC (Fig 4b). Together, these data suggest that suppression of PTPRU in conditions that normally induce quiescence, compromises the ability of hMSC to enter a viable, reversibly arrested state. We infer that PTPRU may sustain cell cycle competence of quiescent hMSC.

**Fig 4.**
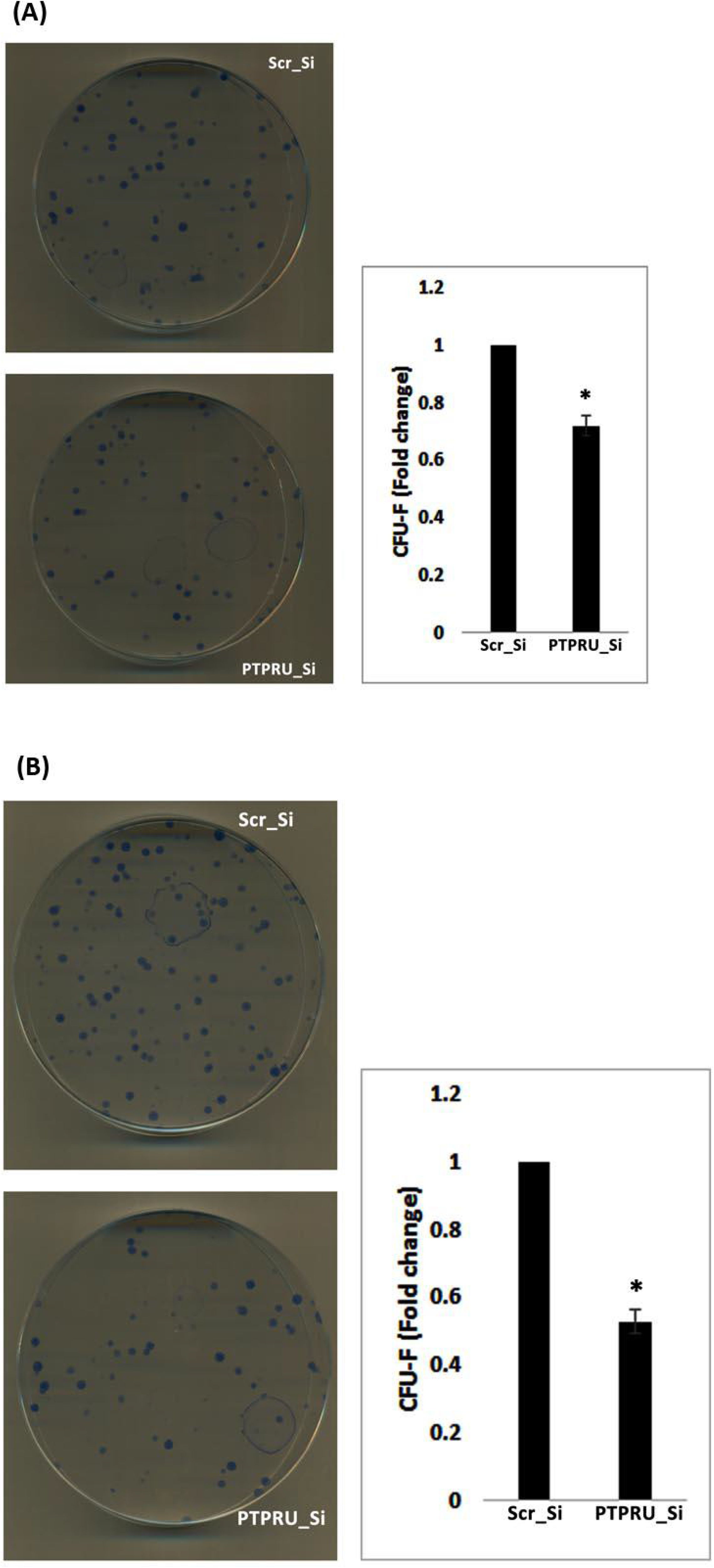
Compromised self-renewal in PTPRU Knockdown MSC. CFU-f assay following PTPRU knockdown in (A) adherent proliferating MSC (B) quiescent MSC. In both A and B, representative images of the stained colonies are shown on the left and quantified on the right. Colonies were counted 10 days after plating at clonal density. Values represent mean + S.D, n=3, p> 0.005.

### Reduction of PTPRU causes up-regulation of osteogenic but not adipogenic lineage genes in hMSC

The failure of PTPRU knockdown cells to exit G0 efficiently suggested the possibility of entry into an alternate pathway of irreversible arrest, which is usually associated with differentiation. To assess whether dampening of proliferative potential in adherent MSC upon PTPRU knockdown is associated with deregulated expression of differentiation genes, we assessed the transcript levels of osteogenic and adipogenic regulators. The expression of osteogenic lineage genes Runx2, Col1A1 and OP was upregulated in PTPRU knockdown MSC in adherent condition as compared to scrambled control siRNA-treated MSC (Fig 5a). However, adipogenic lineage genes did not show a significant change in PTPRU knockdown MSC in adherent condition (Fig 5a). Taken together these findings suggest that normal expression of PTPRU restrains the osteogenic program in adherent hMSC, but does not affect the adipogenic program.

**Fig 5.**
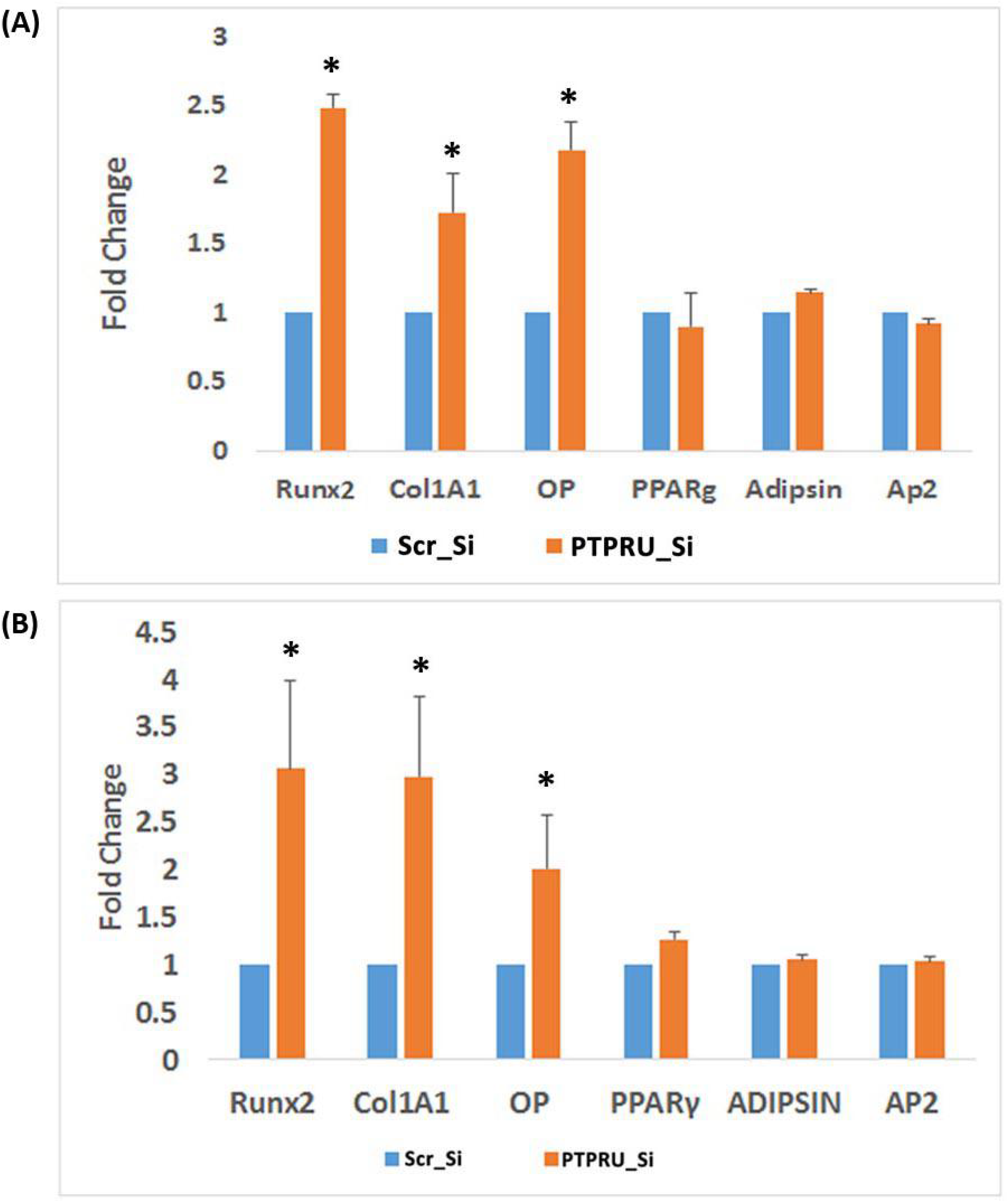
Knockdown of PTPRU leads to deregulated expression of osteogenic but not adipogenic differentiation-associated genes. Transcript levels of osteogenic differentiation genes (*Runx2*, *Col1A1*, *OP*) and adipogenic differentiation genes (*PPAR*γ, *Adipsin*, *ap2*) in PTPRU KD MSC during (A) adherent and (B) suspension culture.

Earlier, we reported that osteogenic but not adipogenic transcripts were induced as hMSC entered G0, which correlated with increased bone differentiation in post-quiescent cultures (Rumman et al. 2018). We therefore assessed whether cell cycle arrest of PTPRU knockdown MSC in suspension also correlates with deregulation of osteogenic genes. qRTPCR analysis of osteogenic and adipogenic lineage genes in suspension-cultured PTPRU knockdown cells showed an increased expression of RUNX2 and Col1A1, whereas no significant change was observed in the expression of PPARγ, AP2 and adipsin (Fig 5b). Thus, suppression of PTPRU in hMSC in adherent conditions is correlated with inhibition of cell proliferation as well as induction of osteogenic lineage genes. Further, PTPRU knockdown in hMSC during suspension condition causes a shift from quiescence-associated reversible arrest to osteogenic differentiation-associated arrest.

Since loss of PTPRU either in adherent or suspension culture resulted in enhanced expression of osteogenic genes, we assessed whether phenotypic differentiation of knockdown cells was altered. Knockdown hMSC (both pre- and post-quiescence) were plated in 96 well plates, and induced to differentiate using osteogenic differentiation medium. The differentiated cells were fixed and stained with Alizarin Red at 5, 10, and 18 days post induction and the extent of osteogenic differentiation quantified. The results showed that PTPRU knockdown led to an early induction of osteogenesis, as evidenced by increased mineralization, in both pre- and post-quiescent cultures compared to control cells (Fig 6a). Taken together, these observations suggest that PTPRU maintains the proliferative potential of hMSC while inhibiting osteogenesis and maintaining the potential for reversible cell cycle arrest. Thus, PTPRU is likely to be involved in a signaling node that balances proliferation, quiescence and differentiation.

**Fig 6.**
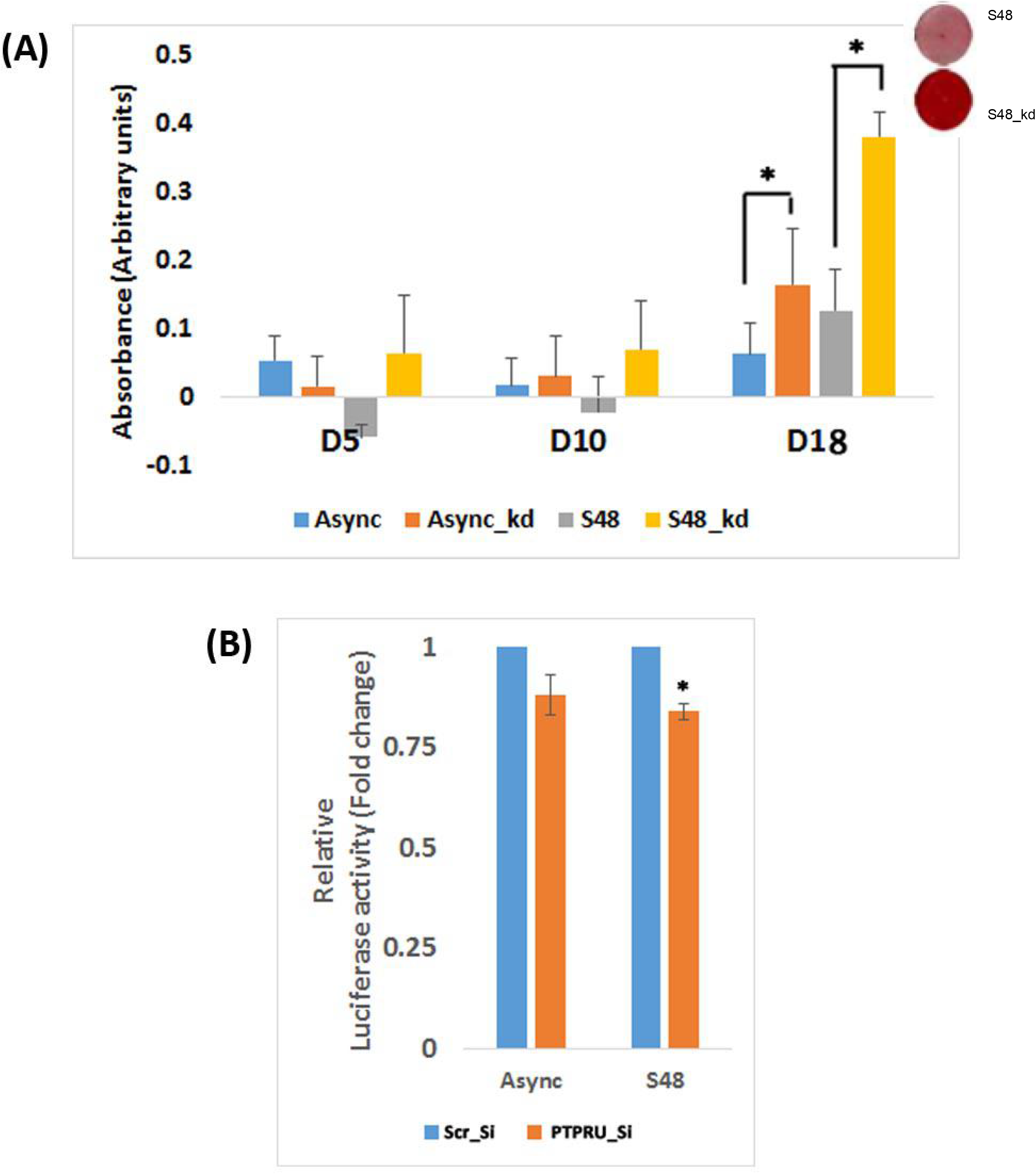
Osteogenic differentiation and TCF-β-catenin transcriptional activity after PTPRU knockdown. (A) Osteogenic differentiation was quantified by estimating calcium deposition using spectrophotometric measurement (A405) of eluted Alizarin Red staining after osteoblast induction at day 5, 10, 18 in proliferating MSC (Async), PTPRU KD proliferating MSC (Async_kd), quiescent MSC (S48) and PTPRU KD quiescent MSC (S48_kd). (B) TCF-β-catenin transcriptional activity assessed by TOP-flash luciferase reporter in control and PTPRU knockdown MSC, in proliferation and quiescence. The inset shows representative images of control and PTPRU knockdown cells stained with Alizarin Red S.

**Fig 7.**
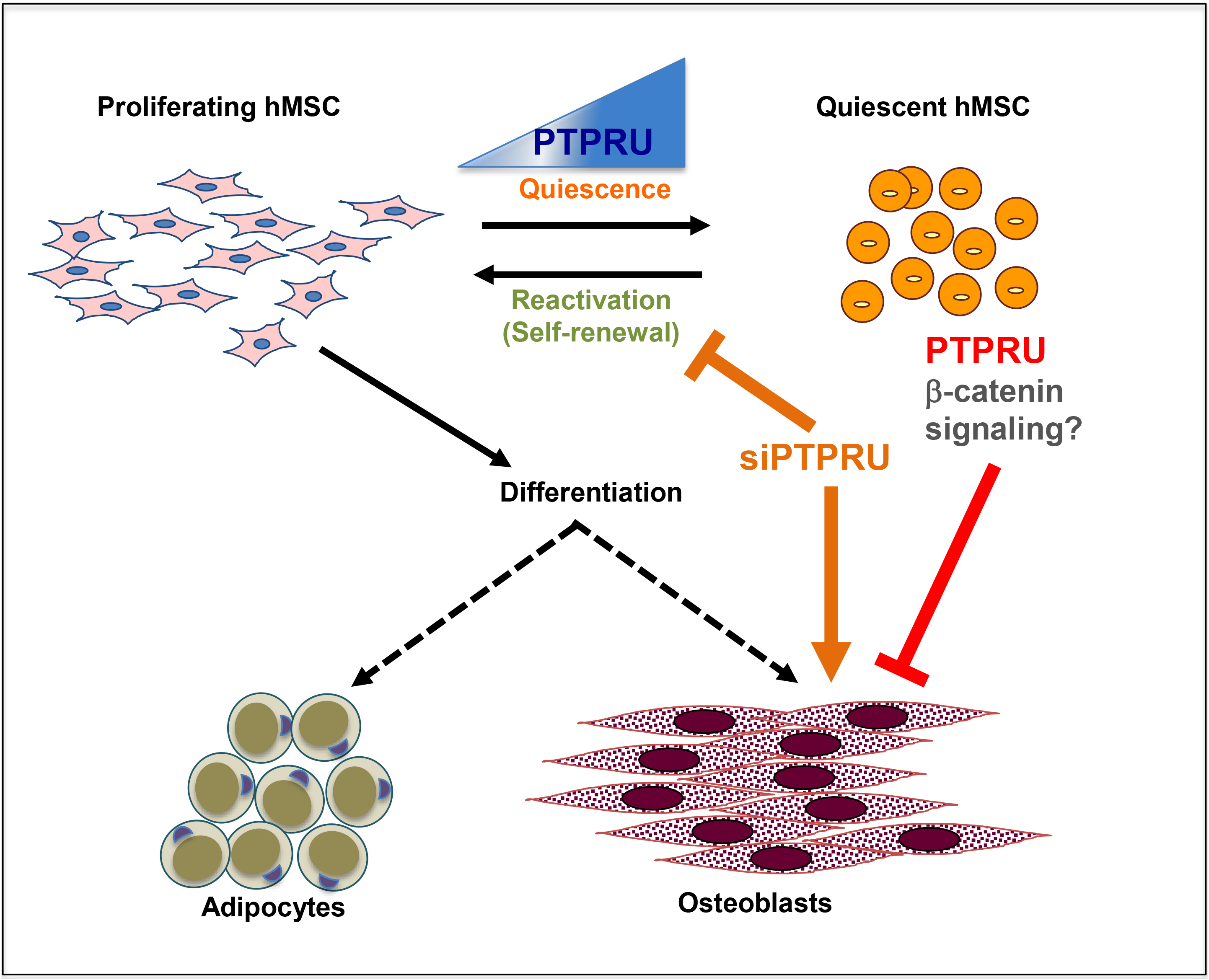
Model for the role of PTPRU in proliferating and quiescent hMSC. In proliferating hMSC, moderate levels of PTPRU maintain proliferation and clonogenicity, potentially via β-catenin. In quiescent hMCS, strong induction of PTPRU suppresses terminal differentiation (particularly osteogenic), thus maintaining the reversibility of the cell cycle arrest. Knockdown of PTPRU leads to mild suppression of β-catenin signaling, correlating with reduced induction of pro-proliferative genes, increased expression of anti-proliferative genes, compromised clonogenic self-renewal, and induction of the osteogenic program.

### PTPRU is a positive regulator of β-catenin activity

Given the key role played by PTPRU in maintaining the quiescence program and mitigating osteogenic conversion in hMSC, we investigated possible mechanisms. Previous studies have showed that full length PTPRU is localized at adherens junctions and binds to β-catenin directly (Yan et al. 2006). This association causes sequestering of β-catenin from the nucleus, thus limiting β-catenin-mediated transcriptional activation. Further, shorter PTPRU isoforms have also been shown to regulate β-catenin transcriptional activity (Zhu et al. 2014).

To assess whether ~75 kDa PTPRU isoform induced in G0 can regulate β-catenin signaling in MSC, we used a TCF reporter assay. Briefly, 12 h post siRNA transfection, hMSC were transfected with the Topflash plasmid (which contains 8 multimerized TCF binding sites upstream of luc reporter) and luciferase activity was measured. For TCF assay in adherent MSC, luciferase activity was measured 48 hours after transfection. To assess luciferase activity in G0, 12 hours after transfection, cells were cultured in methyl cellulose, harvested after 48 hours and reporter activity was measured. We found that knockdown of PTPRU suppresses TCF activity in hMSC, both in adherent and suspended condition (Fig 6b). This observation supports a role of ~75 kDa PTPRU isoform in regulating β-catenin-TCF activity positively.

Taken together, our results show that a quiescence-induced protein tyrosine phosphatase PTPRU promotes reversibility of cell cycle arrest by suppressing osteogenic differentiation with potential involvement of Wnt/TCF signaling.

## Discussion

In this study, we report that a protein tyrosine phosphatase receptor type-U (PTPRU) is specifically induced in reversibly cell cycle arrested hMSCs. Loss of function analysis shows that PTPRU normally sustains self-renewal or cell cycle competence in quiescent hMSCs and suppresses osteogenic differentiation in G0. Further, PTPRU may participate in regulation of Wnt-TCF-βcatenin signaling, a pathway well known to contribute to stem cell function.

Our study supports a context and cell-type dependent role for a 75 kD PTPRU isoform in hMSC. While the reported size of full-length human PTPRU (PTPRU-FL) protein is 200 kD, we find that hMSCs express only a single 75 kD isoform. A single PTPRU mRNA sequence codes for the full-length 200 kD protein (Liu et al. 2014), and no transcript variants have been described. Therefore, it has been hypothesized that shorter isoforms of PTPRU are generated by cleavage of the 200 kD protein, in a manner analogous to other PTP family members, PTPRM & PTPRK, which undergo furin-mediated cleavage to generate shorter fragments (Anders et al. 2006, Burgoyne et al. 2009). Whereas full length PTPRU protein is membrane-bound and is localized at adherens junctions (Yan et al. 2006), the shorter fragments are localized to cytoplasm and nucleus (Zhu et al. 2014).

Several isoforms of PTPRU protein are produced from the single PTPRU transcript in different cell types (Liu et al. 2014). In addition to the 200 kD full-length PTPRU, shorter isoforms of 130 kD, 80 kD and 55 kD have been reported (Zhu et al. 2014). PTPRU-FL is repressed in lung cancer cell lines due to promoter methylation, and restoration of PTPRU-FL expression suppresses proliferation and induces apoptosis (Motiwala et al. 2004). By contrast to the tumor suppressive role of full-length receptor PTPs (RPTP), some of their proteolytic fragments have been shown to contribute to tumorigenesis (Anders et al. 2006, Kaur et al. 2012). For example, fragments of the related RPTPs PTPRK and PTPRM generated through proteolytic cleavage can translocate to the nucleus and activate cell proliferation (Burgoyne et al. 2009), and motility of glioma cells (Kaur et al. 2012). Shorter isoforms of PTPRU are highly expressed in colon cancer, gastric cancer and glioma cell lines (Liu et al. 2014), and shRNA-mediated down-regulation suppresses proliferation and migration (Liu et al. 2014, Zhu et al. 2014). These observations suggest differential roles of full-length and truncated PTPRU proteins in different cancers: while PTPRU-FL shows tumor suppressive functions, shorter PTPRU fragments are oncogenic.

Our study is the first to describe a role for PTPRU in stem cells. Induction of a 75 kDa PTPRU isoform during quiescence suggested a possible role in suppressing proliferation. However, knockdown studies showed that PTPRU is involved in inhibiting hMSC differentiation rather than proliferation. siRNA-mediated knockdown of PTPRU suppressed the expression of the 75 kDa isoform, accompanied by a suppression of proliferation as well as altered expression of cyclins and CDKIs. Moreover, PTPRU knockdown compromises MSC self-renewal ability. This observation may be in consonance with the reported oncogenic role of shorter PTPRU isoforms (Zhu et al. 2014). The dampening of cell cycle in PTPRU knockdown cells was associated with up-regulation of osteogenic lineage differentiation genes, while adipogenic differentiation was unaffected. Thus, in addition to maintaining self-renewing potential in MSC, PTPRU may play a role in lineage choice programs.

Suppression of PTPRU in hMSC did not affect the ability of these cells to exit the cell cycle in non-adherent conditions, but did affect the type of arrest entered by the cells. Moreover, molecular analysis of PTPRU knockdown cells held in G0-inducing conditions revealed that cyclins were hyper-suppressed, whereas expression of CDKIs was upregulated. Further, suppression of PTPRU inhibited reactivation of quiescent cells, even in the presence of adhesion and growth factors. Thus, although PTPRU is normally induced during quiescence, it may not primarily be involved in regulating the cell cycle. Rather, it may be required for other attributes of quiescent cells such as maintaining an un-differentiated state, which can be seen as an uncoupling of cell cycle exit from terminal differentiation. Previously, we had identified Prdm2, an epigenetic regulator of this uncoupling pathway in quiescent muscle cells, and showed that loss of Prdm2 function led to aberrant up-regulation of differentiation lineage genes in quiescence, causing irreversible cell cycle arrest (Cheedipudi et al. 2015). Our current findings in hMSC provide more support for the existence of such uncoupling pathways: we show that osteogenic (but not adipogenic) genes were over-expressed in quiescent PTPRU knockdown cells, correlating with a reduced ability to return to the cell cycle. These observations suggest that PTPRU may help to preserve cell cycle programs in a reversibly repressed state, sustaining the potential for quiescent hMSC to return to active proliferation when required. Taken together, these observations support a role for this RPTP in promoting maintenance of a reversible quiescence program.

Previous studies have shown that PTPs of both receptor type (PTPRF and PTPRQ) as well as non-receptor type (PTPN6) regulate MSC differentiation. PTPRF, PTPRQ, and PTPN6 all negatively regulate adipogenesis (Jung et al. 2009, Kim et al. 2009), while PTPN6 positively regulates MSC osteogenic differentiation at the expense of adipogenesis, via Wnt-β-catenin signaling (Jiang et al. 2016). Although less is known about PTPRU compared to other PTPs, this RPTP has been implicated in control of Wnt signaling. β-catenin is a dual functional protein that plays roles in cell-cell adhesion and transcriptional regulation. In the absence of Wnt ligands, β-catenin is enriched at adherens junctions, its cytoplasmic accumulation held in check by phosphorylation via a GSK3β-Axin destruction complex. Activation of Wnt receptors or inhibition of the destruction complex leads to β-catenin dissociation from adherens junctions, nuclear translocation and transcriptional activation of Wnt-TCF target genes. Wnt-mediated activation of β-catenin function is known to be involved in regulation of quiescence in several adult stem cell types (Rumman et al. 2015, Aloysius et al. 2018).

PTPRU-FL has been shown to interact with and regulate β-catenin signaling through dephosphorylation at Tyr-102 (Yan et al. 2002). Un-phosphorylated β-catenin shows increased interaction with proteins such as E-cadherin, present at plasma membrane (Yan et al. 2006). Smaller PTPRU isoforms have also been shown to regulate β-catenin signaling (Zhu et al. 2014), but whether these shorter fragments possess phosphatase activity is not known. In light of the recent report by Hay et al (Hay et al. 2020), the pseudo-phosphatase domains of PTPRU are not likely to be enzymatically active. Further, unlike PTPRU-FL, the shorter isoforms positively regulate β-catenin transcriptional activity (Liu et al. 2014), by as yet unknown mechanisms including the possibility of acting as dominant negatives.

Our study supports a role for the 75 kD PTPRU isoform in positive regulation of β-catenin signaling in MSC. However two questions remained unanswered: whether the 75 kDa PTPRU fragment interfere with phosphatase activity, and the mechanism by which this short fragment of PTPRU positively regulates β-catenin signaling. It is possible that these shorter fragments are involved in regulating de-phosphorylation of cofactors of β-catenin transcriptional complex, which in turn positively regulate β-catenin signaling. Wnt-signaling through β-catenin is known to be involved in negative regulation of MSC osteogenesis (de Boer et al. 2004, Liu et al. 2011). Our study also supports an inhibitory role for β-catenin signaling in MSC osteogenesis, and a role for 75 kD PTPRU in maintaining a level of β-catenin that dampens bone differentiation.

In conclusion, our results suggest that suppression of PTPRU causes up-regulation of osteogenic lineage genes and reduces self-renewal capability, Thus, PTPRU emerges as a new inhibitor of osteogenic differentiation specifically induced in quiescent hMSC, which may modulate Wnt β-catenin signaling to sustain stem-cell functions in the quiescent state.

## Authors’ contributions

M.R. and J.D. designed experiments, interpreted data and wrote the manuscript; M.R. performed experiments. All authors have read and approved the final manuscript.

## Acknowledgements

We thank Prof. Moustapha Kassem (University Hospital of Odense and the Danish Stem Cell Centre) for useful discussions, and Drs. Prabhavathy Devan (CCMB) and Suchitra Gopinath (THSTI) for critical reading of the manuscript. We gratefully acknowledge CIFF at the Bangalore Life Sciences Cluster for imaging facilities.

